# Dynamic representation of time in brain states

**DOI:** 10.1101/069278

**Authors:** Fernanda Dantas Bueno, Vanessa C. Morita, Raphael Y. de Camargo, Marcelo B. Reyes, Marcelo S. Caetano, André M. Cravo

## Abstract

The ability to process time on the scale of milliseconds and seconds is essential for behaviour. A growing number of studies have started to focus on brain dynamics as a mechanism for temporal encoding. Although there is growing evidence in favour of this view from computational and *in vitro* studies, there is still a lack of results from experiments in humans. We show that high-dimensional brain states revealed by multivariate pattern analysis of human EEG are correlated to temporal judgements. First, we show that, as participants estimate temporal intervals, the spatiotemporal dynamics of their brain activity are consistent across trials. Second, we present evidence that these dynamics exhibit properties of temporal perception, such as the scalar property. Lastly, we show that it is possible to predict temporal judgements based on brain states. These results show how scalp recordings can reveal the spatiotemporal dynamics of human brain activity related to temporal processing.

## Introduction

Humans and non-human animals are able to estimate temporal intervals across a wide range of scales^1,2^. Intervals ranging from hundreds of milliseconds to seconds are specially critical for sensory and motor processing, learning, and cognition. On this scale, the theoretical dominant models are based on the existence of an internal clock, consisting of a pulse emitting oscillator and an accumulator that counts the pulses^3–5^. Electrophysiological evidence for this model consists mainly in ramping activity as a possible correlate of the accumulation of pulses, and has been found using invasive recordings in brain regions as prefrontal, parietal and motor areas. In human electroencephalography (EEG), it has been suggested that the contingent negative variation (CNV), a slow cortical potential of developing negative polarity could be a reflect of such processes, given its similarity with the hypothesized characteristics of an accumulation process^6^. However, in both cases (invasive recordings and human EEG) it is not clear whether ramping activity is coding time or using temporal information to anticipate or react to events^7^. Moreover, the existence of an internal clock has been criticised based on behavioural findings^8^, and for being biologically unrealistic.

For this reason, a number of alternate models of timing, which take into account neural data, have been put forward as biologically-plausible explanatory candidates, such as state-dependent networks models^9^. For this class of models, neural circuits would be inherently capable of temporal processing as a result of the complexity of cortical networks coupled with the presence of time-dependent neuronal properties^8^. In this view, neural systems can take advantage of the temporal evolution of neural states, caused by the variation in neural and synaptic properties of the system. When activated, a neural system would follow a unique trajectory through its state space. Thus, by adapting networks to read out specific neural states they could be tuned towards specific temporal intervals^9^. However, the vast majority of evidence in favour of these models rely on computational modelling or *in vitro* studies^8, 10–12^ and evidence from human and non-human animal recordings are still sparse^13,14^.

One of the main difficulties of investigating this class of models using non-invasive human electrophysiological recordings (EEG) is how to characterise different states and their temporal evolution. On the other hand, recent EEG studies have suggested that it is possible to differentiate spatially overlapping brain states by analysing participant-specific patterns^15–19^. This methodology has been successfully applied to magneto-encephalographic recordings and was able to dissociate between standard and deviant tones, frequent versus rare melodies, visual stimulus location and stimulus orientation^15^. In the present study, we analysed the time-resolved EEG signals using multivariate pattern analysis (MVPA), to investigate whether the evolution of brain states over time can carry information about the temporal interval tracked by participants and also their future behavioural responses.

## Results

Human participants (*n* = 14) performed a temporal categorisation task. The experiment was based on a “shoot the target” task that has been previously used to study temporal perception^20, 21^. The task consisted of two types of trials. In Regular trials, a visual target transited from the periphery of the screen towards a central target zone taking 1.5 seconds to reach screen centre, where there was an “aiming sight” (Figure 1a). In these trials, participants were instructed to press a button in order to produce a “shot” (an audiovisual stimuli) when the target passed the aiming sight. In Test trials, the trajectory of the target was masked by an occluder and only the aiming sight remained visible throughout the entire trial. In these trials, participants did not see the movement of the target, but only heard the start of its movement by a characteristic sound that targets made when they started moving (both in Regular and in Test trials). One automatic shot was given in each Test trial in one of seven different intervals (0.8,0.98,1.22,1.5,1.85,2.27 or 2.8 s) after the sound indicated that the target started moving. Participants had to judge whether the shot occurred at an interval “shorter”, “equal”, or “longer” than the time the target normally took to reach the screen centre. Test and Regular trials were presented in a pseudo randomized order during the experimental session in which three consecutive Test trials were not allowed.

**Figure 1.**
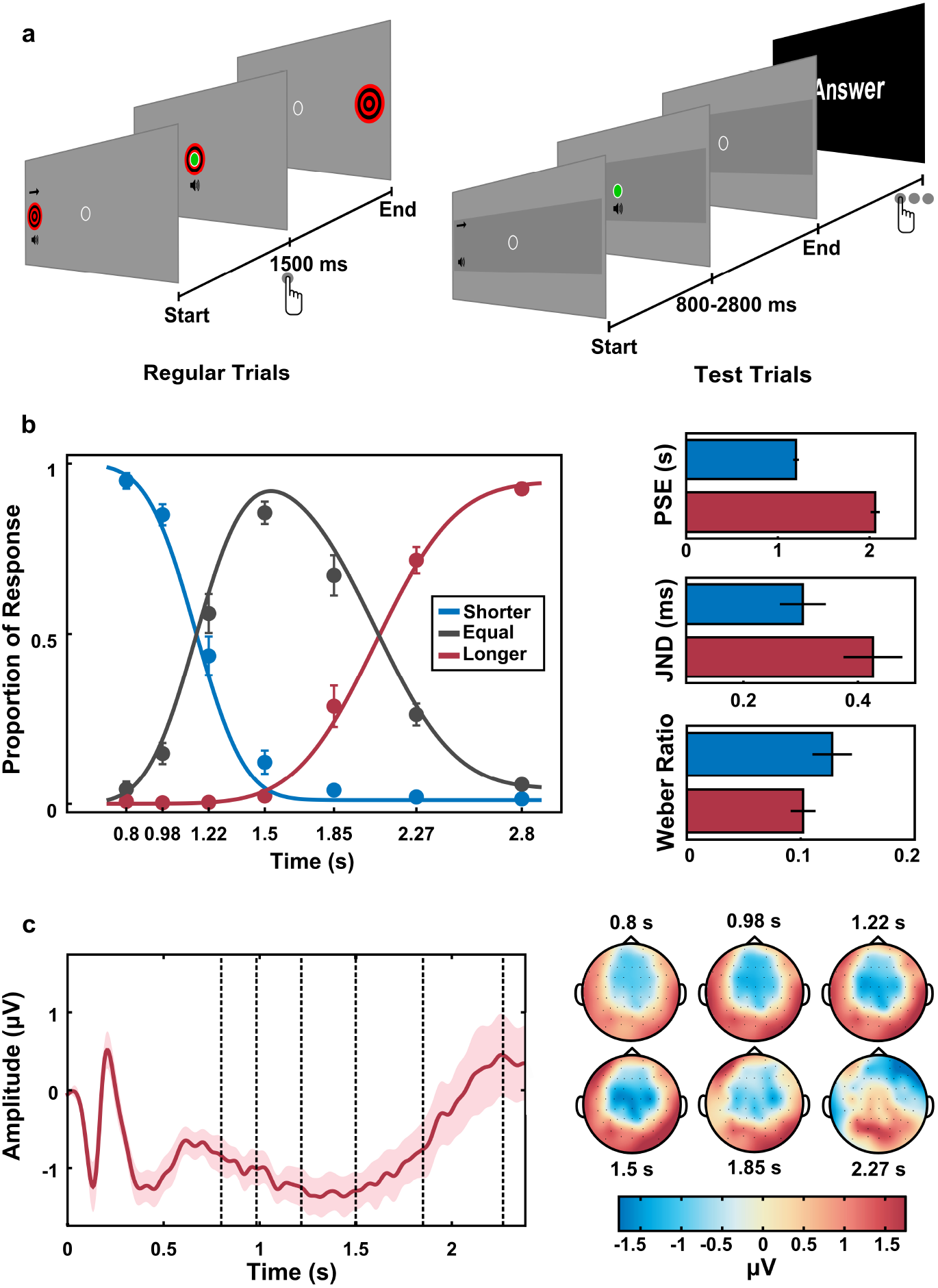
Experimental task and results. (a) The task consisted of a computerised shoot the target task. In regular trials a bulls-eye moved towards the centre of the screen reaching it in 1.50 seconds. Participants were instructed to produce a “shot” when the target passed trough the “aiming sight” fix in the centre of the screen. In Test trials, target trajectory was masked and automatic shots were given at seven different intervals (0.80,0.98,1.22,1.50,1.85,2.27 and 2.80 seconds). Participants had to judge whether the shot occurred after an interval “shorter”, “equal”, or “longer” than the time the target normally took to reach screen centre. (b) Psychometric functions describing performance on Test trials. Right panel shows estimated parameters from the psychometric functions: Points of Subjective Equality (PSE), Just Noticeable Difference and Weber ratio. Plots show mean and standard error of the mean (s.e.m.) across participants. (c) Contingent Negative Variation (CNV, mean ± s.e.m.) for central-parietal electrodes for Test trials longer or equal to 2.27 seconds (dashed lines indicates intervals where the second marker could have been presented). The CNV peaks at the memorised interval. Right panel shows the topographies for the intervals close to possible moments of target presentation.

The proportion of each response was calculated for each interval and two independent cumulative normal functions were fitted to the proportion of responses “shorter” and “longer”. As expected, participants made few errors when categorising intervals that were clearly shorter, equal or longer than 1.5 seconds (Figure 1b). Participants demonstrated highest uncertainty (point of subjective equality, PSE) between shorter and equal responses for intervals around 1.20 ± 0.03 s (mean ± s.e.m) and between equal and longer responses for intervals around 2.10 ± 0.05 s (paired t-test between estimated PSEs, *t*_13_ = 15.05, *p* < 0.001, Bayes Factor (BF) in favour of the alternative> 100).

A hallmark of interval timing is its scalar property (a generalisation of the Weber Law), which states that the variability in temporal estimations increases linearly with the magnitude of the interval estimated. That is, errors in temporal estimations scale with the durations of the intervals^3^, which implies that shorter intervals are easier to discriminate than longer intervals. Accordingly, in our experiment, participants were more sensitive (i.e., responded differentially) to small temporal differences across shorter intervals compared to longer intervals. This is illustrated by the smaller Just Noticeable Difference (JND) for the shorter compared to the longer intervals (*JND_short_* = 0.287 ± 0.031 s, *JND_long_* = 0.438 ± 0.032 s, paired t-test, *t*_13_ = 4.77, *p* < 0.001, BF in favour of the alternative= 77.49). As predicted by the scalar property, when sensitivity was normalised by interval length no difference between shorter and longer intervals was observed (*Weber_short_* = 0.118 ± 0.013, *Weber_long_* = 0.103 ± 0.006, paired t-test *t*_13_ = −1.39, *p* = 0.187, BF in favour of the null= 1.64).

For the EEG recordings, we focused our analysis on trials in which the interval was at least 2.27 s long, collapsing the data from the two longest intervals. The event-related potential for central electrodes can be seen in Figure 1c. Consistent with previous results, there was a clear Contingent Negative Variation potential (CNV, a slow cortical potential of developing negative polarity) which peaked at the reference interval^6, 7^.

For a neural system to be able to read time by its trajectory through state space, the trajectory of the activity elicited by a target must be consistent across activations. Thus, we checked if the recorded dynamics were consistent across trials. Indeed, the pattern of the EEG signals across the scalp followed a structured sequence in time during the different trials (cluster-stat=2349, *cluster* – *p* < 0.001, Figure 2a). Next, we used the Mahalanobis distance^19, 22^ to perform pairwise comparisons across time points to determine whether the pattern of the EEG signal contained information about the interval between events. As shown in Figure 2b, multivariate distances between time points followed a diagonal-shaped pattern (i.e., a stronger similarity across points closer in time), suggesting a sequential activation of overlapping states.

**Figure 2.**
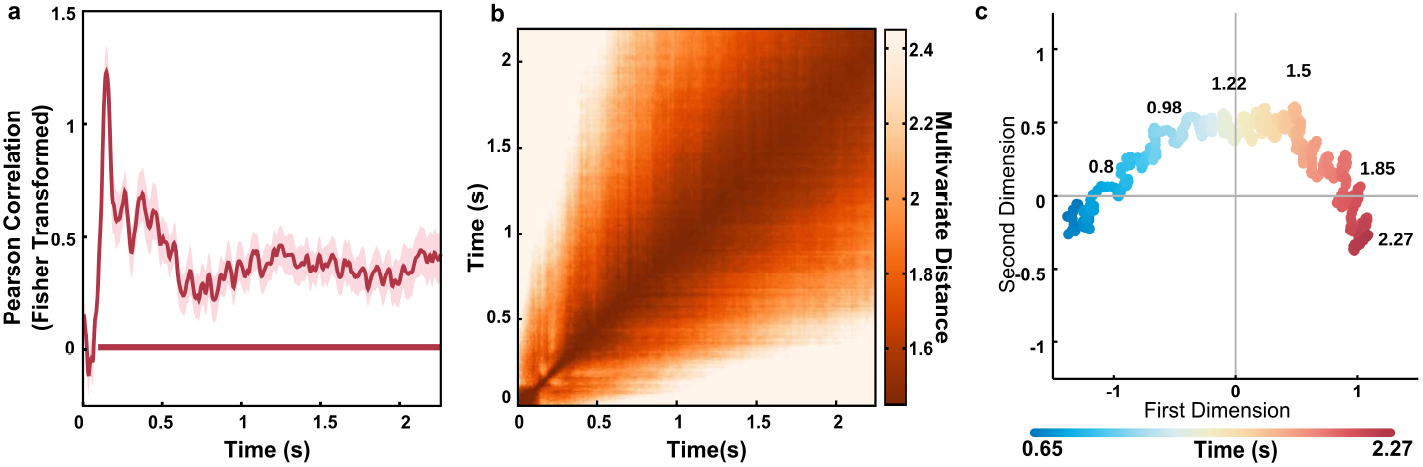
Spatiotemporal dynamics and temporal perception (a) Similarity index of the spatiotemporal dynamics across trials (mean ± s.e.m). Red lines at the bottom represent the temporal cluster where similarity was significant. (b) Pairwise multivariate distance matrix between all time points. There is a strong similarity for time points closer in time, suggesting a sequence of activation states. (c) Multidimensional distance between the activity in different time points for the period between 0.65 s and 2.2 s after the first cue was presented visualised in two dimensions using multidimensional scaling (MDS). The colour of each point represents its physical interval. The trajectory represents the path linking the sequence of activation states.

The evolution of the spatiotemporal dynamics captured by the EEG sensors can be visualised using multidimensional scaling and plotted against the first two dimensions. We focused on the period starting 0.65 s after the presentation of the first sound to minimise the influence of the event related potentials. This trajectory represents the path linking the sequence of activation states, while the multidimensional distance between time points in state space reflects the difference in the overall response captured by all sensors. In line with the pairwise distances showing similarity in activations for points closer in time, the recovered trajectory preserves the temporal information of the activated states (Figure 2c). There is a strong correlation between distance in time and distance in state space.

To quantify the relation between state space and behaviour, we focused our analysis at time points when the interval could have ended. We performed multivariate pairwise comparisons (using Mahalanobis distances) on data for the six first intervals (0.8,0.98,1.22,1.5,1.85,2.27 seconds) and used multidimensional scaling to represent them in a two dimensional plot (Figure 3a).

**Figure 3.**
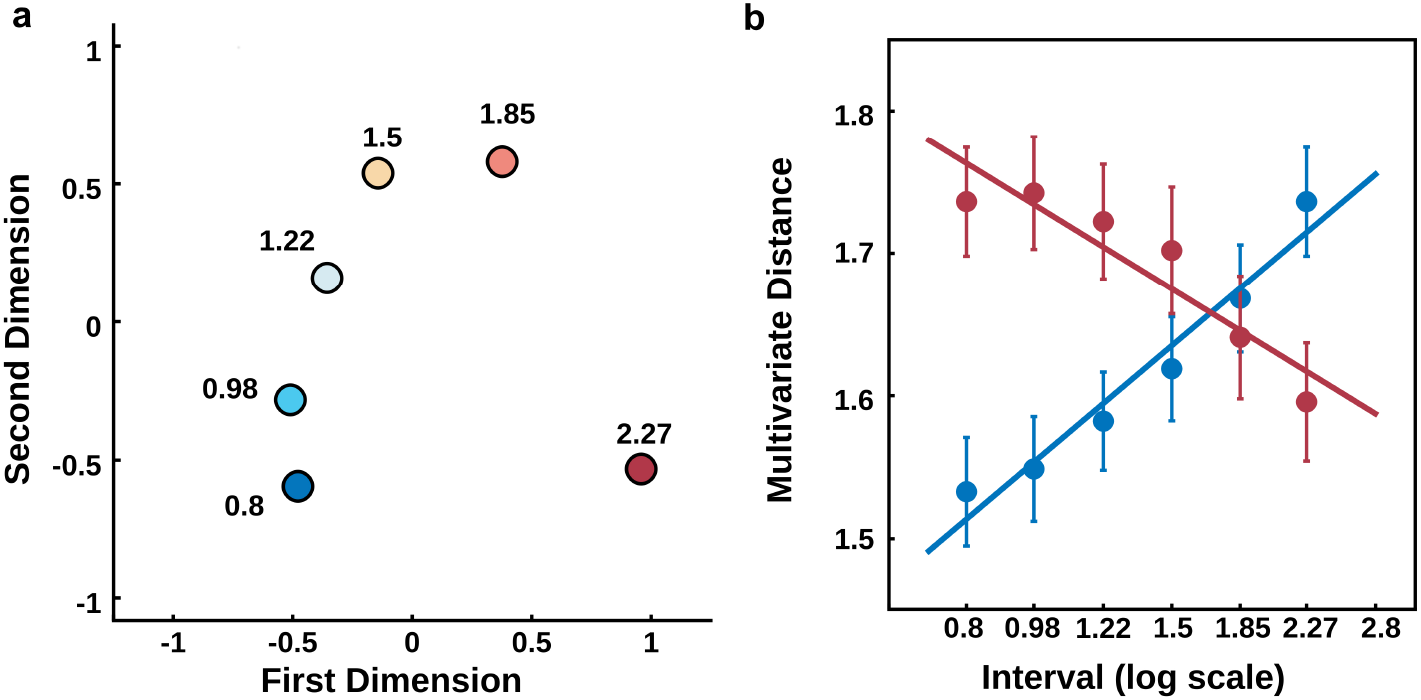
Distance in state space is correlated to distance in time (a) Multidimensional distance between the activity in the different possible intervals visualised in two dimensions using multidimensional scaling (MDS). The colour of each point represents its physical interval. (b) Mean distances in state space (Mahalanobis distance) as a function of temporal separation (*log*_10_ scale). Blue (red) markers shows pairwise multivariate distances (mean ± s.e.m) between the 0.8 s (2.27 s) and all other intervals. The slope of the fitted linear functions indicated that the rate of change in state space as a function of time is faster for the first than for the last interval.

To quantify the relationship between distance in time and in state space, we fitted linear functions to the log-transformed temporal distance with the state space distance, comparing the first and last interval to all others. As shown in Figure 3b, there was a strong association between both distances, suggesting that states further apart in time are also further apart in state space (fitted slope for shortest interval = 1.58±0.08, t-test to zero:*t*_13_ = 19.48, *p* = 0.001, BF in favour of the alternative> 100, fitted slope for longest interval = −1.29 ± 0.09, t-test to zero: *t*_13_ = −13.50, *p* = 0.001, BF in favour of the alternative> 100). Importantly, the slope for the shortest interval is steeper than for the longest interval (paired t-test on the absolute estimated slopes, *t*_13_ = 3.348, *p* = 0.005, BF in favour of the alternative= 9.68). This suggests that, for the shortest interval, the rate of change in state space as a function of time is higher than for the last interval, once again in accordance with the scalar property of time.

Thus far, we have shown that the properties of brain states revealed by MVPA follow key characteristics of temporal processing. However, if they are related to temporal perception, then it should be possible to decode participants' temporal judgements based on the recovered states. To test this possibility, we focused our analysis on single trial data from the two intervals closest to when participants had highest uncertainty about their temporal judgement: (1) 1.22 s, for which participants had a high uncertainty on whether it was shorter or equal to 1.5 s; and (2) 1.85 s, for which participants had a high uncertainty on whether it was equal or longer than 1.5 s.

When participants had a high uncertainty between shorter/equal responses, positions within the path of state space that were closer to the equal state increased the probability of equal responses (t-test on the estimated slopes, *t*_12_ = 2.28, *p* = 0.041, BF = 1.84 against the null, Figure 4b). In a related analysis, we compared the location in state space where participants were when they responded shorter or equal. We found that equal responses were associated with positions closer to the equal state (*t*_12_ = 2.20, *p* = 0.048, BF = 1.66 against the null, Figure 4c).

**Figure 4.**
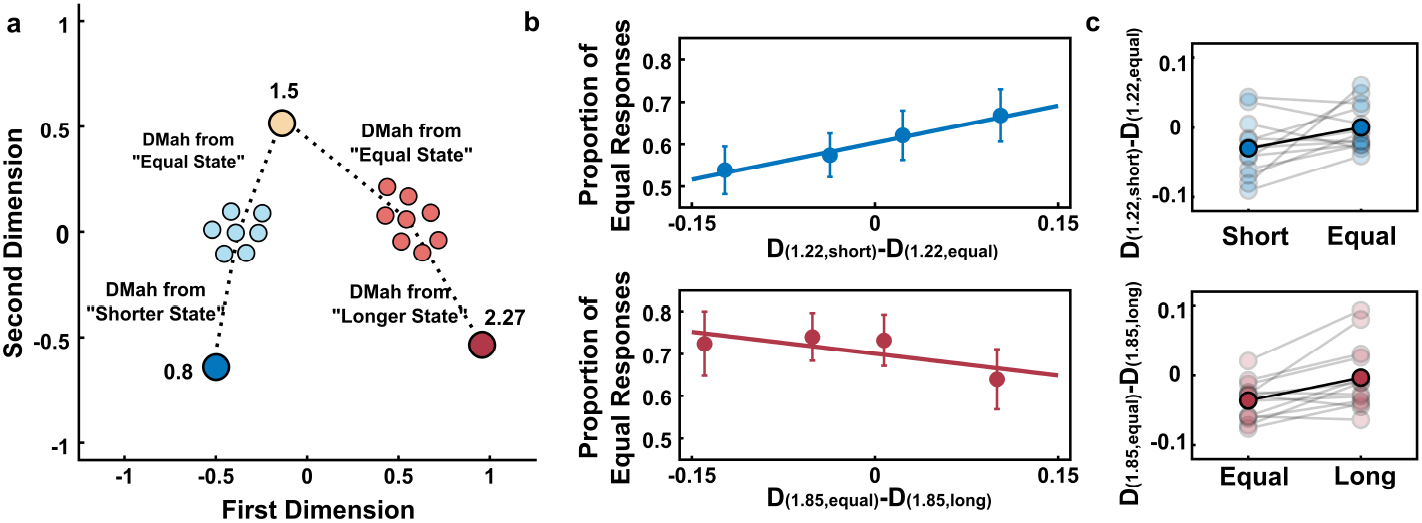
Correlation between position in state space and behaviour (a) Schematic representation of the methodology used to estimate position in state space for single trials. These analyses focused in intervals where participants had maximum uncertainty on whether the interval was shorter than or equal to 1.5 s (1.22 s) and equal or longer than 1.5 s (1.85 s). For each of these trials, multivariate distances between activity in that trial and two other landmarks were estimated. The difference between these distances was used as an index of position in state space. (b) Proportion of Equal responses as a function of position in state space. Each dot represents a quartile split of “equal” responses (mean ± s.e.m) as a function of the position where participants are in state space. The continuous line represents the fitted function. For shorter intervals, proportion of Equal responses increased as activity was more similar to the equal landmark (upper panel). For longer intervals, proportion of Equal responses decreased when activity was more similar to the longer landmark (lower panel). (c) Position in state space as a function of the given response. Upper panel shows mean position in state space when participants responded “shorter” or “equal” for trials with a 1.22 s interval. Lighter circles and lines show data across participants and darker circles and lines show the mean across participants. Negative (positive) values indicate that activity is more similar to the shorter (equal) state. Lower panel shows mean position in state space when participants responded “equal” or “longer” for trials with a 1.85 s interval. Negative (positive) values indicate that activity is more similar to the equal (longer) state.

A similar pattern was found when participants had a high uncertainty between equal/longer responses. Positions within the path of state space that were further from the equal state decreased the probability of equal responses (t-test on the estimated slopes, *t*_12_– = −2.81, *p* = 0.016, BF = 3.87 against the null). We further compared the location in state space when participants responded longer or equal. We found that equal responses were associated with positions closer to the equal state (*t*_12_ = 3.41, *p* = 0.048, BF = 9.20 against the null, Figure 4c).

Within each case, the physical interval tested was identical in all trials, showing how trial-by-trial fluctuations in the distance travelled in state space can partially account for the variability in temporal judgements.

## Discussion

In this study, we investigated whether time-resolved EEG signals can carry information about time. We have shown how dynamic pattern analyses can characterise states correlated with coding of temporal intervals. These states show properties classically related to temporal perception, such as scale invariance, and were predictive of temporal judgements.

A growing number of studies in non-human animals have investigated if the neural mechanisms of temporal perception in the range of hundreds of milliseconds to seconds are related to population dynamics instead of a single central clock^1,13,23^. According to this view, temporal information emerges from neural properties that are naturally time-varying, such as short-term synaptic plasticity^24^. Our results show, for the first time to our knowledge, how these dynamics can be captured by non-invasive electrophysiological recordings, addressing an important lack of evidence in favour of this population dynamics view in humans.

The majority of timing studies in humans have focused on the contingent negative variation (CNV) as a potential signature of the subjective experience of time^6^. However, the exact role of the CNV on temporal perception has been recently criticised both on experimental and theoretical terms^7,25^. For example, a recent study has argued that post-interval evoked potentials (e.g., activity evoked by a stimulus that marks the end of an interval) can reflect the subjective experience of time better than the CNV^25^.

Our results, on the other hand, suggest that temporal experience can be tracked during the interval to be estimated. Although the averaged amplitude of the CNV in a group of electrodes might not be a good predictor of the elapsed interval, other subtle differences present in individual electrodes, and possibly hindered by the CNV, could actually carry information about time. When a network is activated it will follow a complex trajectory dependent not only on the synapses directly activated by the input, but also on the ongoing activity of the network^9^. This process leads to distinct spatiotemporal patterns of activity, which, according to our results, produce different patterns of activation across the EEG sensors. This finding adds to the increasing evidence of how multivariate methods represent a powerful approach to decode task-relevant dimensions.

As these trajectories evolve through time in a consistent way, stimuli presented in different time points will find the network in a different state leading to different activity patterns^9^. Therefore, although small differences in the state of a network might not be detectable using classical EEG methods, they could still lead to marked differences of evoked activity by new inputs in different time points, as recently reported^25^.

Although our results are consistent with the state-dependent timing models, it could be argued that our analyses are capturing properties of other models of temporal processing, as central clock models. For example, the recovered states could be related to the accumulator process: as pulses are accumulated, they generate activity that is reflected across electrodes. However, it is important to mention that, in a broad view, state-dependent models can be extended to be consistent with the majority of the timing models^26^, with different models imposing specific constraints on what would define the state space. Thus, although our findings open an important and missing first step in how to investigate state dependent models in human EEG, they do not, at the moment, present conclusive evidence in favour of such view.

One important question that remains to be answered is whether these recovered brain states are part of a coding scheme used to track time or a by-product of other processes that could generate a time-decodable signal. For example, in our specific task, the activity decoded could be biased due to particular strategies adopted by participants, as imagining the movement of the target to estimate time. Future experiments with different tasks could easily test such possibility. Moreover, comparing how well these decoding methods work on different tasks can help to determine whether these methods are capturing a more automatic or cognitively controlled system for temporal processing^27,28^.

A more general formulation of the same argument can be proposed. One could argue that the majority of neural processing has a temporal structure. If this structure is consistent enough, even if not directly used to track time, it could lead to similar results reported by our study. Once again, future studies can address this issue by observing how those brain states behave in tasks that have similar temporal structures, but different temporal demands.

Taken together, our results show an important proof of principle that the analyses of multivariate patterns can bring important advances in our understanding of temporal processing. Using a simple unsupervised learning methodology, we have shown that activity captured across all electrodes carry information that is correlated to temporal perception and to subsequent temporal judgements. Given the absence of clear electrophysiological correlates of temporal processing in human EEG, we propose that the use of multivariate pattern analyses can be of extreme importance, specially in situations where different theoretical models make similar behavioural predictions. There is a myriad of temporal tasks and illusions that can be used to study time in humans. Future studies can combine these tasks and use similar methods herein described to elucidate how temporal information can be encoded at the population level to support time-dependent functions.

## Methods

### Participants

Sixteen volunteers (age range, 22 – 27 years; 9 female) gave informed consent to participate in this study. The number of participants was chosen based on previous experiments. All of them had normal or corrected-to-normal vision and were free from psychological or neurological diseases. The experimental protocol was approved by The Research Ethics Committee of the Federal University of ABC. Two participants were excluded from the analyses because more than 3 channels during EEG recordings presented an increased number of artifacts.

### Stimuli and Procedures

The experiment consisted of a computerised “shoot the target” task^20^. The stimuli were presented using Psychtoolbox *v*.3.0 package for MATLAB on a 17–inch CRT monitor with a vertical refresh rate of 60 Hz, placed 50 cm in front of the participant. Responses were collected via a response box with 9 buttons (DirectIN High Speed Button; Empirisoft). Each trial started with the presentation of a target (1.5 visual degrees radius, red and black) at the left hemifield of the screen (background RGB-colour 150; 150; 150) and an “aiming sight” (an empty circle with 0.5 visual degree radius) at the centre of the screen. After a random interval (500 ms-1000 ms) a beep (1000 Hz, 70 dB, 100 ms duration) was presented simultaneously with the start of movement of the target from left to right. The target moved at a constant speed (9 degrees/sec) taking 1.50 seconds to reach the centre of the screen, thus passing through the aiming sight. In Regular trials (350 trials) a button press (with the right index finger) produced a “shot” (a green disc, presented inside the aiming sight) and a second beep (500 Hz, 70 dB, 100 ms duration), presented simultaneously. Participants were instructed to hit the target by pressing the button at the appropriate moment.

Test trials followed the same schema, but the trajectory of the target was masked by a gray rectangle (3 visual degrees of height, RGB-colour 130; 130; 130) and automatic shots were given at seven different intervals (0.8,0.98,1.22,1.5,1.85,2.27 or 2.8 s). At the end of each Test trial, participants had to judge whether the shot occurred at an interval “shorter”, “equal”, or “longer” than the time the target normally took to reach the screen centre. Participants responded using three different buttons. Responses in Test trials were unspeeded and could be given starting 800 ms after the automatic shot. Trials were presented in a random order, with the restriction of having a maximum of three Test trials in sequence. Participants performed a total of 10 blocks, each with 35 Regular and 35 Test trials. The first 10 trials in each block were always Regular trials. The experimental session lasted 60 minutes on average.

### EEG recordings and pre-processing

EEG was recorded continuously from 64 ActiCap Electrodes (Brain Products) at 1000 Hz by a QuickAmp amplifier (Brain Products). All sites were referenced to FCz and grounded to AFz. The electrodes were positioned according to the International 10 – 10 system. Additional bipolar electrodes registered the electrooculogram (EOG). EEG pre-processing was carried out using BrainVision Analyzer (Brain Products). All data were down-sampled to 250 Hz, re-referenced to the average of all electrodes, filtered (0.05 Hz to 30 Hz), epoched from 500 ms before the first beep to 1000 ms after the second beep and baselined from −150 ms to 0 ms relative to the first beep. An independent component analysis (ICA) was performed to reject eye movement artifacts. Eye related components were identified by comparing individual ICA components with EOG channels and by visual inspection. The number of trials rejected for each participant was small (13% on average). ERP analysis were performed on data using the SPM8 and Fieldtrip toolboxes for MATLAB. The CNV for Figure 1c was estimated at central-parietal electrodes (C3,C1,Cz,C2,C4).

### Behavioural Analysis

Behavioural analysis was based on the proportions of each type of response as a function of interval. For each participant, two independent sigmoidal functions were fitted to the data. The first function was fitted on the proportion of “shorter” responses and the second on the proportion of “longer” responses. The psychometric data from each participant and condition were fitted with Cumulative Normal functions, each defined by four parameters: threshold, slope, lapse-rate and guess rate^29^. Guess rates were fixed at 0 across all participants and conditions, and lapse rates were restricted to a maximum of 0.05. The three free parameters were fitted separately for each participant. The points of highest uncertainty were estimated as the predicted interval corresponding to 50% of “shorter” responses and the predicted interval corresponding to 50% of “longer” responses. Quality of fit for each participant was assessed by correlating predicted values to observed responses (r-square 0.96 ± 0.01; lowest individual r-square = 0.88). At the group level, parameters were compared using paired t-tests and Bayes factors in favour of the null or alternative hypothesis are reported^30,31^.

The parameters for the sigmoidal functions were fitted using maximum likelihood estimation as implemented in the Palamedes Toolbox^32^. To measure temporal sensitivity for shorter and longer intervals, we calculated Just Noticeable Difference (JND) scores, which represents the absolute difference in seconds between the intervals at which 25% and 75% of shorter or longer responses were given. Weber fractions were estimated as:

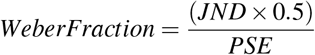

### Multivariate Pattern Analysis

#### Similarity between different trials

To calculate the similarity of the sequence of patterns activated we used a bootstrap approach. For each participant, data from the two longest trials were collapsed, divided in two, and averaged across trials within them. Then, data from each set and time point (two row vectors with the averaged amplitude values of all 62 electrodes) were compared using a Pearson correlation (Rho’). This procedure was repeated 5000 times for each participant and Fisher transformed coefficients were averaged across permutations for each participant.

At the group level, Fisher transformed Rhos’ were compared to zero using a paired t-test. Statistics of one-dimensional EEG-analyses were inferred non-parametrically^19, 33^ with sign-permutation tests. For each time-point, the Fisher transformed Rhos' of each participant was randomly multiplied by 1 or −1. The resulting distribution was used to calculate the p-value of the null-hypothesis that the mean Rhos value were equal to 0. Cluster-based permutation tests were then used to correct for multiple comparisons across time using 10,000 permutations, with a cluster-forming threshold of p < 0.05. The significance threshold was set at p < 0.05 and all tests were two-sided.

#### Dissimilarity Matrix

To determine whether the pattern of the EEG signal across channels contained information about the elapsed time, we used the Mahalanobis distance to perform pair-wise comparisons between different time points. Several studies have shown that the Mahalanobis distance is superior to Euclidean distance because it accounts for the covariance structure of the noise between features^19,34^. A similar bootstrap approach as in the above analyses was used. For each participant, data from the two longest intervals (2.27 and 2.8) were collapsed, randomly split in two halves and averaged within each half. A pairwise multivariate dissimilarity (Mahalanobis distance) of each time point from one half to all others time points in the other half were calculated as follows:

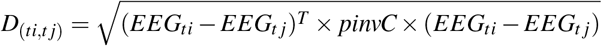

where *EEG_ti_* and *EEG_tj_* are row vectors containing the average signals of the two time points being compared from the two independent halves. The *pinvC* is the pseudo inverse of the error covariance matrix, estimated by pooling over the covariances of both time points being compared, estimated from all trials, using a shrinkage estimator that is more robust than the sample covariance for data sets with many variables and/or few observations^19, 34^. This procedure was performed for all time points, from 0 to 2.27 seconds, in 4 ms bins. Pairwise multivariate dissimilarity matrices were estimated separately for each participant and iteration. This procedure was repeated 2500 times for each participant and the mean dissimilarity matrix was calculated for each participant. The grand mean dissimilarity matrix was calculated by averaging individual matrices across participants.

#### Multidimensional Scaling

Trajectory in state space was visualised using metric multidimensional scaling (MDS) as implemented in MATLAB. In Figure 2c, MDS was performed on the grand mean pairwise dissimilarity for time points between 0.65 s and 2.27 s matrix and data was plotted against the first two dimensions (*stress* < .001). For Figure 3a, MDS was performed on the grand mean pairwise dissimilarity for possible intervals and data was plotted against the first two dimensions (*stress* < .001)

#### Correlation between dissimilarity and performance

To calculate how single trial trajectories correlated with performance, we focused on the two intervals closest to the points of highest uncertainty (1.22 s and 1.85 s). To measure where in the state space trajectory participants were in each trial, we compared single-trial states to two other landmarks states, allowing us to estimate the location in the state space where participants were when the interval ended. The landmarks consisted of the averaged data from pre-stimuli period (−100 to 0, relative to target onset) for the 0.8 s interval (short landmark), 1.5 s interval (equal landmark) and 2.27 s (longer landmark). Importantly, these landmarks were estimated on the averaged data of the two longest intervals (collapsing 2.27 s e 2.8 s, as done previously). Thus, the single-trial data and the data used to estimate the landmarks were completely independent.

For the 1.22 s, data from the last 100 ms before target presentation (1.12 – 1.22 s) was averaged, resulting in a row vector for each trial (with each row containing data from one electrode). This was then compared to the short landmark and the equal landmark, using the Mahalanobis distance as previously described. The subtraction *D*_(1.22, *short*)_ – *D*_(1.22, *equal*)_ resulted in an index of how similar the state in that particular trial was to each landmark. Trials where the state is more similar to the equal landmark yields higher values. If these distances are correlated to performance, then the proportion of equal responses should increase as this index increases. To test this hypothesis, we performed two separate, although related, analyses: (1) the estimated indexes were used as a predictor in a generalised linear model regression, with a probit link for the binomial distribution. This procedure was performed separately for each participant and at the group level the estimated slope coefficients were tested against zero using a paired t-test. (2) the position for “equal” and “shorter” responses were averaged for each participant and compared at the group level using a paired t-test.

Similarly, we compared single-trial data for 1.85 s to equal and longer landmarks and used the difference *D*_(185, *equal*)_ – *D*_(185 *longer*)_. In this case, higher values indicate a stronger similarity to the longer state and should be correlated with a decrease in the proportion of equal responses. The same analyses as described above were performed.

In both cases, data from one participant was removed due to a small proportion of one of the response types (< 10%).

## Acknowledgements

This work was supported by the Fundação de Amparo à Pesquisa do Estado de São Paulo (FAPESP) research grants 15/00794-2, 15/04554-6 and 13/24889-7 and by Conselho Nacional de Pesquisa (CNPq) research grant 122942/2014-0. The authors would like to thank Dean Buonomano, Nicholas E. Myers and the members of the Timing and Cognition Laboratory at UFABC (http://neuro.ufabc.edu.br/timing/) for useful discussions and suggestions on earlier versions of this manuscript

## Author contributions statement

A.M.C., M.S.C., F.D.B and V.C.M. conceived the experiment. F.D.B. and V.C.M. performed the experiments. All authors analysed the data, wrote and reviewed the manuscript.

## Additional information

Competing financial interests: The authors declare no competing financial interests

